# Computational design of Bax-inhibiting peptides

**DOI:** 10.1101/2024.10.28.617283

**Authors:** Tom Vlaar, Bernadette Mayer, Lars van der Heide, Ioana M. Ilie

## Abstract

The proteins of the Bcl-2 family play crucial roles in regulating apoptosis. It is divided into pro-survival and pro-apoptotic proteins that determine cellular fate. In particular, Bax is a crucial executor of apoptosis as its activation initiates the apoptotic phenotype. Hence, targeting this protein represents an attractive therapeutic approach, which can aid in regulating apoptotic signalling and potentially contribute to the development of novel therapies against cancer and neurodegenerative diseases. Here, we introduce a digital paradigm, which relies on rational design and computer simulations to develop and validate peptide-based agents that bind to Bax, thereby inhibiting its apoptotic properties. The peptides are rationally designed and optimized to bind to Bax starting from the crystal structures of affimers in complex with Bcl-2 proteins. Next, molecular dynamics simulations (MD) are employed to probe the stability of the Bax-peptide complexes and to estimate the binding free energies. The results show that the designed peptides bind with high affinity to Bax. Two of the designed peptides bind in the canonical hydrophobic groove (BH1 domain) of Bax and one peptide binds to the outside of the BH3 domain (*α*_2_-helix). Notably, the peptides restrict the flexibility of the *α*_1_-*α*_2_ loop, modulating the bottom trigger site associated with toxicity. All in all, the results highlight the potential of these peptides as valuable tools for further exploration in modulating apoptotic pathways and set the structural foundation for a machine learning powered engine for peptide design.

## Introduction

Programmed cell death or apoptosis is a tightly regulated process in multicellular organisms.^1^ Dysregulation of this mechanism can lead to several diseases, including cancer and possibly neurodegenerative disorders such as Parkinson’s disease. ^2,3^ In the healthy cell, the B-cell lymphoma 2 (Bcl-2) protein family maintains the homeostasis of the mitochondrial apoptotic pathway through complex interactions,^4,5^ while also regulating mitochondrial dynamics, the endoplasmatic reticulum, calcium storage and autophagy. ^6^ The Bcl-2 family consists out of 26 currently known members that are classified into pro-survival (or anti-apoptotic) Bcl-2 proteins, signalling pro-apoptotic members (or BH3-only proteins) and executor proteins.^7^ The pro-survival proteins inhibit cell death by binding to the pro-apoptotic Bcl-2 proteins and vice-versa. In response to cellular stress BH3-only proteins are activated which in turn activate the executor proteins. The executor proteins transfer from the cytosol to the mitochondrial outer membrane where they accumulate, oligomerize, and facilitate mitochondrial outer membrane permeabilization releasing cytochrome c and other factors.^8,9^ In dopamine neurons Mcl-1 is a critical Bcl-2 pro-survival factor as its chemical inhibition has been shown to activate the pro-apoptotic protein Bax, caspases and result in neuronal cell death.^3^ As the loss of dopamine neurons is a hallmark of Parkinson’s disease Mcl-1 function may be related to disease onset and possibly provide a therapeutic target.^3,10^

The Bcl-2-associated X protein (Bax) is a proapoptotic Bcl-2 family protein, which shares structural and sequence similarities with other Bcl-2 family proteins, such as the anti-apoptotic Mcl-1.^11^ Bax consists of nine *α*-helices, which are structured around a hydrophobic core composed of helices *α*_2_-*α*_5_ and has a globular structure (Fig. 1 - left panel). In its inactive form, the trans membrane *α*_9_-helix is folded into the hydrophobic groove. In the active state, the helix is inserted into the mitochondrial membrane. It has been proposed that the *α*-helices of the BH3 domains of Bim (BH3-only protein) can transiently bind to an activator site near the N-terminus (*α*_1_/*α*_6_) of Bax, thereby causing its activation.^12^ This interaction displaces the *α*_9_-helix from the hydrophobic groove and initiates mitochondrial outer membrane integration. The opening of the hydrophobic groove provides the possibility of interaction by the activator BH3 domains, further inducing conformational changes to Bax where unfolding of the *α*_2_ helix occurs followed by the dissociation of both *α*_1_ and the *α*_6_-*α*_8_ latch, and forming homodimers with neighbouring molecules by inserting the everted BH3 domain into their hydrophobic groove.^13,14^ Another activation binding site was reported at the proximal *α*_1_-*α*_2_ loop in mitochondrial Bax triggered by generated monoclonal antibodies, opening up new possibilities for activation of Bax besides the use of BH3-only proteins.^15^ The vMIA protein (viral mitochondria localized inhibitor of apoptosis) binds Bax at the *α*_3_-*α*_4_ and *α*_5_-*α*_6_ hairpins and was shown to have inhibitory effects.^16^ Adjacent to this binding site is the Bcl-2 (pro-survival protein) BH4 domain binding domain, consisting of residues located on *α*_1_, the *α*_1_-*α*_2_ loop, the *α*_3_-*α*_4_ and *α*_5_-*α*_6_ hairpins,^17^ which contribute to the inhibition of Bax mediated apoptosis by restricting conformational changes. Small molecule binding at the same site allosterically activates Bax. ^18^

**Figure 1:**
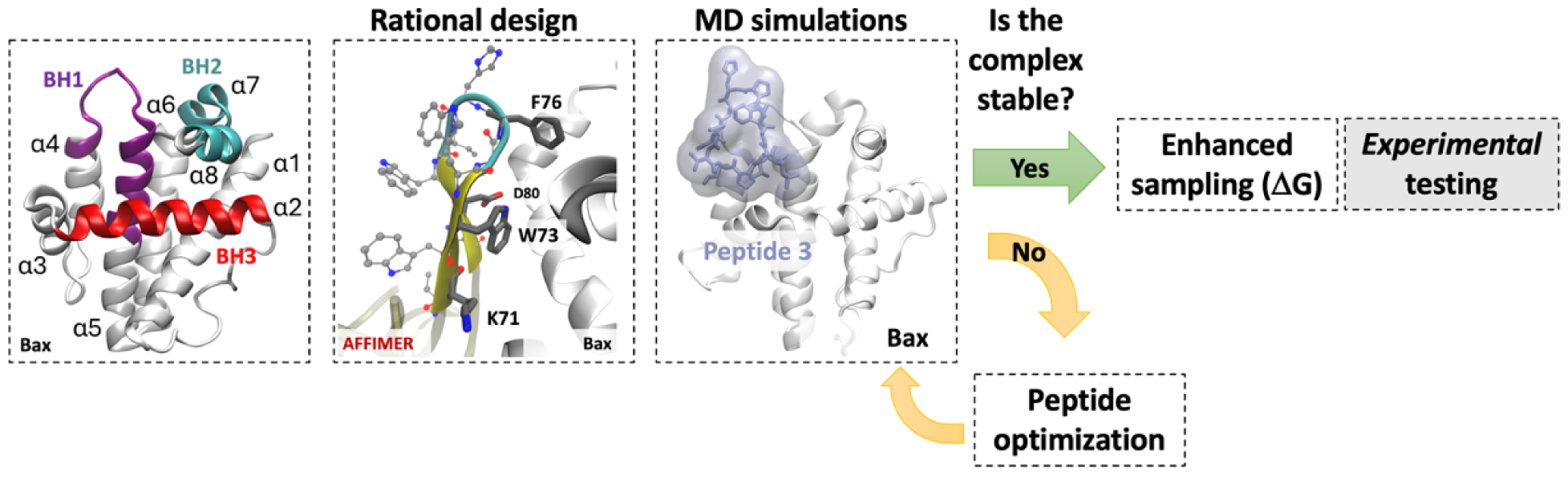
Peptide development strategy. Bax protein in its inactive state (PDB ID: 1F16^11^). Here, the BH1 domain is coloured in purple, BH2 domain in orange, BH3 domain in cyan and the *α*_9_-helix in red (left panel). The peptides are rationally designed from available 3D structures of protein complexes. Briefly, the residues that interact with the protein are grafted from an affimer. The terminal residues are then cyclized via head-to-tail cyclization and the complexes are subjected to molecular dynamics simulations. The peptides that do not stably attached to Bax are optimized via single point mutations. If the complex is stable, the binding affinities of the peptides to Bax are determined and the best candidates are advanced into experimental testing.

Cyclic peptides are rapidly evolving as therapeutics and are emerging as powerful in-hibitors in the drug development field. ^19,20^ Cyclic peptides are developed to combine conformational rigidity and solubility to enable binding to undruggable interfaces with high affinity.^21^ Cyclic peptides have proven to be excellent candidates for cancer therapy, ^22^ organ transplantation^23^ and inhibition of amyloid aggregation.^24,25^ Their size and functional properties ensure that the contact area is large enough to provide high selectivity, their ability to form salt-bridges and hydrogen bonds can lead to strong binding affinities,^26^ and cyclization increases their proteolytic stability. ^27^ Engineering new peptides with tailored properties and high affinities towards a desired targets is a non-trivial, resource-demanding and challenging task. Experimentally, phage display or mRNA display allow the generation of large libraries of peptides with target specificity.^28^ These libraries can produce a vast array of peptides, but the chemical synthesis and the numerous experimental trials require significant resources.

Digital design and simulations are complementary tools that help to overcome some of these difficulties. For instance, recent advances with digital tools like RosettaFold^29^ and AlphaFold2^30,31^ can help with the determination of three-dimensional (3D) high resolution structures of protein-peptide complexes. These conformations can then be used as starting structures for molecular dynamics simulations to probe stability and dynamics, which can then be leveraged to design better binders potentially using machine learning techniques prior to experimental testing.^19^ The advantages are three-fold. First, the simulations have atomistic resolution and can provide information on the dynamics of the complex and the isolated peptides, which exceed experimental resolution.^32^ Second, the simulations allow the exploration of a vast parameter space, which can help in the optimization of the designed peptides. Third, this step reduces the number of experimental trials to be carried out and increases the number of potent binders that can be generated and designed.

Recently, we proposed a recipe for generating a digital twin that would rely on information from computational and experimental findings to simulate the effect of a cyclic peptide-based drug on amyloidogenic targets.^19^ This digital twin would therefore require the efficient incorporation of data from different sources, including binding constants, conformations, specificity etc., to enhance the design and optimization of future peptide-based drugs. Here, we take the first step towards building this digital twin and we use Bax as a model system. Hence, we introduce a novel strategy to design cyclic peptide-based binders that can compete with Mcl-1 for binding to Bax, thereby freeing up Mcl-1 and enhancing cellular resilience.^6,11^ For this, we introduce a novel digital strategy that relies on rational design and molecular dynamics simulations to develop and validate the new binders *in silico* prior to experimental validation (Fig. 1). First, we rationally design cyclic peptides starting from known three dimensional structures of BCL-2 family members in complex with non-antibody scaffold proteins. Second, we optimize the peptide sequences via single point mutations to enhance binding to the target. Third, we probe their structural stability and estimate their binding free energies to Bax by using (enhanced sampling) molecular dynamics simulations. The results reveal the mechanisms of interaction between three optimized cyclic peptides and Bax and characterize their binding to the target. Furthermore, the calculated binding free energies show that the peptides favorably bind to Bax. This aids in understanding the mechanisms behind cyclic peptide-Bax stability and provides the starting information for building a digital twin tailored for cyclic peptide design.

## Theory and Methods

### System preparation

The aim of the present study is to develop novel peptide-based ligands that compete against MCL-1 against Bax binding. For this, the 15-166 segment of the pro-apoptotic protein Bax was extracted from solution NMR (PDB: 1F16^11^), Fig. 1 - left panel. To reduce computational cost, the disordered N-terminal residues 1-14 were removed. Additionally, residues 167-192, forming the *α*_9_-helix that mediates the formation and bioactivity of heterodimers, were removed to simulate Bax in its active state. For the rational design of the cyclic peptides, the X-ray diffration structures of MCL-1 and BCL-xL in complex with affimers were used as scaffold (PDB IDs: 6STJ and 6HJL,^33^ respectively).

### Rational design of cyclic peptides

Each system was prepared by aligning the complexes of MCL-1 or BCL-xL and affimers with the homologous Bax structure. The peptides were defined starting from the interfacial residues for the affimers, independently. In particular, the residues located at the epitopes and the amino acids in the loops of the affimer sequences that point towards the proteins and contribute to the binding affinities were isolated. ^33^ Subsequently, each peptide was optimized via single point mutatations, head-to-tail cyclized, and the covalent bond only was minimized using Maestro 2023-3.^34^

Following this protocol, three peptides were designed. Peptide ^37^QGGVNPEEM^45^ (P1) was grafted from the chain E affimer residues sourced from the MCL-1-affimer complex (PDB: 6STJ^33^). To avoid steric clashes, residue M38 was mutated to G38. The negative control (NC) ^37^QKKGGGEER^45^ is derived from P1 by introducing the G38K, G39K, V40G, N41G, P42G and R45M mutations to disfavor binding. Peptide ^68^VWVKRDLVFGGPENFK^83^ (P2) was designed from the first loop of the affimer chain D sourced from the BCL-xL-affimer structure (PDB: 6ST2^33^). Peptide ^70^VKPALLWSPHGNF^82^ (P3) was engineered from the first loop of the affimer chain C, extracted from the BCL-xL-affimer structure (PDB: 6HJL^33^). The topology of the new Bax-cyclic peptide system was then subjected to long molecular dynamics simulations to investigate the stability of the complex.

### Simulation protocol

Four sets of simulations were carried out using the same simulation parameters. First, unrestrained 1 *µ*s MD simulations were conducted in duplicate in the NVT ensemble. For the systems that proved stable, *i.e.,* no peptide detachment or sliding/reattachment of the peptide at secondary locations, a stable complex conformation was selected as the starting configuration for the next set of simulations. For the systems that were not stable, *i.e.,* the peptide detached from the surface of Bax, single point mutations in the peptides were introduces to avoid steric clashes (e.g. P1) and the protocol was repeated. Second, ten 500-ns simulations of the selected complexes were run to probe the statistical stability of the complex and characterize the peptide-Bax interactions. Third, ten 500-ns simulations of Bax and the peptide alone were performed to compare the structural stability and properties of the protein and the peptides in the bound and unbound states. The bound starting configuration was selected from the simulations of the complex. Fourth, umbrella sampling simulations were performed to determine the binding free energies of the peptides to Bax.

#### Simulation details

The simulations were carried out using the GROMACS 2020.4 simulation package. All simulations were performed using the all-atom CHARMM36m force field^35,36^ and the TIP3P water model.^37^ The systems were solvated in cubic boxes with edge lengths of 7 nm and 4.3 nm for the Bax and peptide only systems (3.7 nm for P3 only), respectively. The Bax-peptide complexes were solvated in cubic boxes with edge lengths of 7.3 nm (8.2 nm for the Bax-P2 complex). Each system was neutralized and a background concentration of 150 mM of NaCl was added. Steepest decent energy minimisation was followed by a two-step isothermal-isobaric ensemble (NPT) equilibration. The temperature and pressure were kept constant at of 300 K and 1 bar using the velocity rescaling^38^ (modified Berendsen) thermostat and Berendsen barostat,^39^ respectively. The temperature and pressure coupling times were fixed to 0.1 and 2 ps, respectively. The NPT equilibration was performed in two steps with position restraints of 1000 kJ·mol^-1^·nm^-2^ on the heavy atoms for 5 ns, followed by an equilibration with restraints of 100 kJ·mol^-1^·nm^-2^ for 5 ns to gently equilibrate the newly generated protein-peptide complex. The production simulations were performed in the NVT ensemble in absence of restraints. The short range interaction was cutoff beyond a distance of 1.2 nm and the potential smoothly decays to zero using the Verlet cutoff scheme. For the energy and pressure, a long-range dispersion correction was applied. To compute long-range electrostatic interactions, the Particle Mesh Ewald technique ^40,41^ was employed with a cubic interpolation order, a real space cutoff of 1.2 nm, and a 0.16 nm grid spacing. A fourth order LINCS algorithm with two iterations was employed to constrain all bond lengths.^42^

#### Umbrella sampling

To determine the intermolecular binding energies between the peptide and Bax, umbrella sampling was used.^43,44^ In essence, an additional energy term, a bias, is applied to the system to ensure efficient sampling along a chosen reaction coordinate to connect energetically separated states. Here, the distance between the centers of mass of the peptide and the protein, *d*, was used as the reaction coordinate. The reaction coordinate of range *d* is then divided into several “windows” centered at values *d_i_* where the harmonic bias potential *ω_i_*(*d*) only restricts the reaction coordinate in the *i* ^th^ window to fluctuate around *d_i_* by:^45^

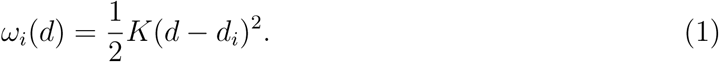

with *K* the force constant. This changes the total energy of the system to *E*^b^(*i*) = *E*^u^(*i*) + *ω_i_*(*d*) with *E* being the total energy and the superscripts ‘b’ and ‘u’ denoting the biased and unbiased quantities, respectively. The unbiased free energy for window *i*, *A_i_*(*d*), is based on the probability distributions 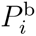 and the bias potential and can be obtained by

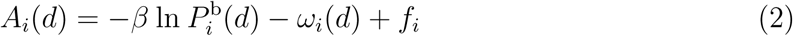

where *β* = 1*/k_B_T*, *k_B_* being the Boltzmann constant, *T* the temperature and, *f_i_* a window-dependent offset: *f_i_* = −(1*/β*) ln⟨*exp*[−*βω_i_*(*d*)]⟩. The windows are then combined using the weighted histogram analysis method (WHAM)^46–48^ to determine *f_i_*.

To generate the series of configurations along the reaction coordinate, chosen as the center of mass distance between the peptide and the protein, the peptide was pulled away from the protein along the z-axis. As starting configuration, a stable conformation of the ten 500 ns simulations was used. The peptides were pulled with a pull-rate of 0.01 nm/ps over the course of 400 ps of MD using a harmonic potential with a force constant of 1000 kJ·mol^-1^·nm^-2^, saving snapshots every 1 ps. In addition, the backbone of the protein was restrained with a force constant of 1000 kJ·mol^-1^·nm^-2^. The windows were sampled along the z-axis with a spacing of 0.2 nm and a force constant of 1000 kJ·mol^-1^·nm^-2^ for Bax-P1 complex. For both P2- and P3-Bax complexes a spacing of 0.1 nm, a force constant of 5000 kJ·mol^-1^·nm^-2^ were used. For the umbrella sampling simulations, the box was elongated along the direction of the pull by 5 nm and the same parameters were used for the MD simulations of the windows. Each umbrella window was simulated in the NPT ensemble for 305 ns (except for P1, which reached convergence after 105 ns) from which the first 5 ns were considered to be part of the equilibration.

## Results and discussion

### Peptide binding reduces Bax flexibility

For each Bax-peptide complex, two initial simulations of 1 *µ*s were performed to investigate the structural stability. The results from these sets of simulations revealed that the negative control detaches from the surface of Bax, and does not reattach at secondary locations. Peptides P1, P2 and P3 slide on the surface of Bax within the first few ns of the simulations and converge to new interaction hotspots, which they preserve throughout the simulations. Specifically, P1 rearranges and binds at the *α*_2_-helix in the BH3 domain. Both P2 and P3 attached to the BH1 domain, comprised of mainly the *α*_4_-*α*_5_ loop and part of the *α*_5_-helix (Fig. 2(a)). These conformations were the selected as initial configurations for ten 500-ns simulations to investigate complex stability and peptide binding effects on the structure and dynamics of Bax.

**Figure 2:**
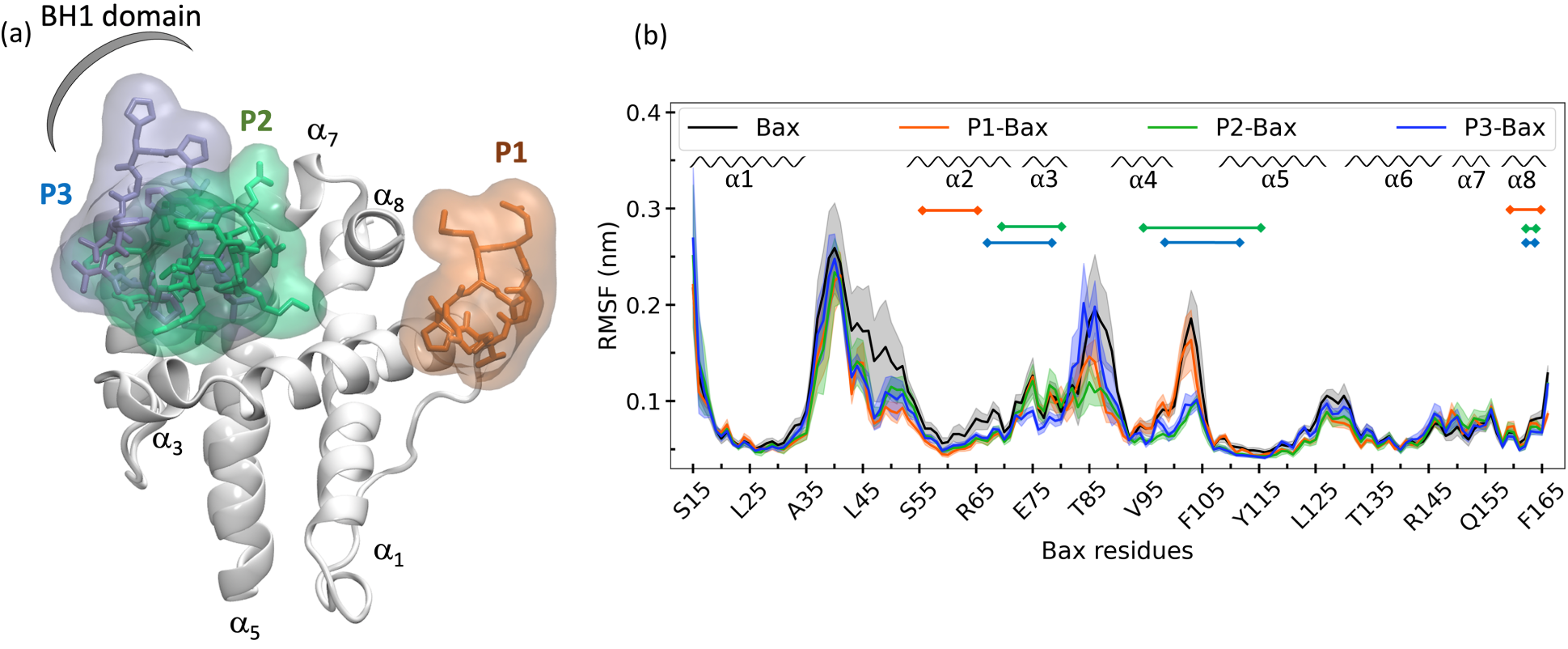
Structural overlap of Bax in complex with the peptides. Highlighted are P1, P2 and P3 in orange, green and blue (a). Root mean square fluctuations (RMSF) profiles of Bax (b). Compared are the free Bax (black), Bax in complex with P1 (orange), Bax in complex with P2 (green) and Bax in complex with P3 (blue). The RMSF profiles are calculated as the average over 50 independent 100-ns profiles. The shaded areas represent the standard error calculated as the standard deviation of the RMSFs of the 50 independent 100-ns profiles. For the P1 simulations only 30 independent 100-ns profiles are used from the systems with a stable interface. Highlighted are the secondary structure elements of Bax (top, and the binding sites of the peptides (horizontal lines).

The analysis focused on the structural stability of the complex, reveals that P2 and P3 remain attached to Bax in all the 5-*µ*s simulations, with average deviations of from the reattached structures of the three bound peptides 0.2 ± 0.1 nm and 0.17 ±0.02 nm, respectively (Fig. S1). P1 is the least stable as it detaches in four of the ten simulations and occasionally reattaches at a secondary interaction site. Hence, only the runs, in which P1 remains stably attached (average deviations of 0.27 ±0.02 nm, Fig. S1) to Bax are considered for the analysis. In absence of peptides, Bax is structurally stable with average deviations from the crystal structure of 0.22 ± 0.03 nm. In complex with the peptides the average deviations from the crystal structure are reduced (0.19 ±0.01 nm, 0.16 ±0.02 nm and 0.17 ±0.01 nm in the Bax-P1, Bax-P2 and Bax-P3 complexes, respectively, Fig. S2), suggesting that the peptides modulate the stability of Bax.

The analysis focused on the protein flexibility shows that peptide attachment has marginal impact on the plasticity of the secondary structure elements of Bax and affects to a higher degree the fluctuations of the loops (Fig. 1(b)). Specifically, low fluctuations of the secondary structure elements are observed independently of the free or the complexed Bax state, except for *α*_2_, which is decreased upon peptide binding. In contrast, peptide attachment reduces the mobility of the *α*_1_-*α*_2_ and *α*_4_-*α*_5_ loops. The flexibility of loop *α*_3_-*α*_4_ is differently modulated by the three peptides *i.e.,* P3 has no impact, while P1 and P2 reduce its plasticity, with the latter having a more pronounced effect. This modulating effect in the loops can be linked to the binding sites of the peptides. P1 binds at the C-terminus of the BH3 domain, which contributes to the reduced flexibility of the *α*_2_-helix and the allosteric stiffening at the loops. P2 and P3 contact the *α*_4_-*α*_5_ loop, which leads to the stabilization of the BH1 domain, comprised of mainly of this loop and part of the *α*_5_-helix Furthermore, during the simulations P2 was was observed to enter the canonical hydrophobic groove (*α*_2_-*α*_5_ ^49^) and form multiple contacts at both sites of the groove (to be discussed in the following paragraphs). By entering the pocket, the groove opens, thereby reducing the flexibility of the connecting *α*_3_-*α*_4_ loop. P3, which also binds more superficially in the hydrophobic groove at the BH1 domain as compared to P2. As a result, the plasticity of the *α*_3_-*α*_4_ loop is reduced. Furthermore, the flexibility of the *α*_3_-*α*_4_ remains comparable to the one of Bax in the free state but restricts the motion of *α*_3_ as compared to the other systems. The subtle differences in the effects of P2 and P3 may be ascribed to the longer sequence in case of P2 (sixteen residues compared to thirteen residues P3), which enables a larger contact area with the protein.

### Peptide binding modulates the opening of the hydrophobic groove and closes the trigger bottom pocket

To further investigate the modulating effects of the peptides on the hydrophobic groove, the structural and dynamic details of the *α*_3_-*α*_4_ domain were analyzed (Fig. 3(a)). The impact on the flexibility of the loop reflects in the solvent accessible surface area (SASA) of the *α*_3_-*α*_4_ sequence (spanning residues M74-M99) (Fig. 3(b)). Specifically, the binding of P1 and P2 leads to an increase in SASA, with the latter having a more pronounced effect. This correlates with the reduced flexibility of the loop (Fig. 2(b)), despite the peptides binding at different sites on the surface of Bax. In contrast, P3, which does not affect the flexibility of the loop, reduces the solvent accessibility of the canonical hydrophobic groove. The peptide binding effect is also reflected in specific distances between the helices, *i.e.,* M74-M99 and A81-R89 highlighting the *top* and the *bottom* of the groove, respectively (Fig. 3(c-d)). Hence, the reduced flexibility upon P1 attachment translates into a rearrangement of the helices characterized by the opening of the groove at the top and closing at the bottom (*scissor* motion). The closing of the loop is also associated with the unfolding of the C-terminus of the *α*_3_-helix, which populates more coil structures (Fig. S4). P2 attachment leads to an increase of about 0.2 nm in both the top and the bottom distances of the groove, which is a consequence of the insertion of the peptide in the canonical hydrophobic groove. Thus, the peptide P2 induces the opening of the groove, whereas P1 allosterically opens the top and closes the bottom of the groove.

**Figure 3:**
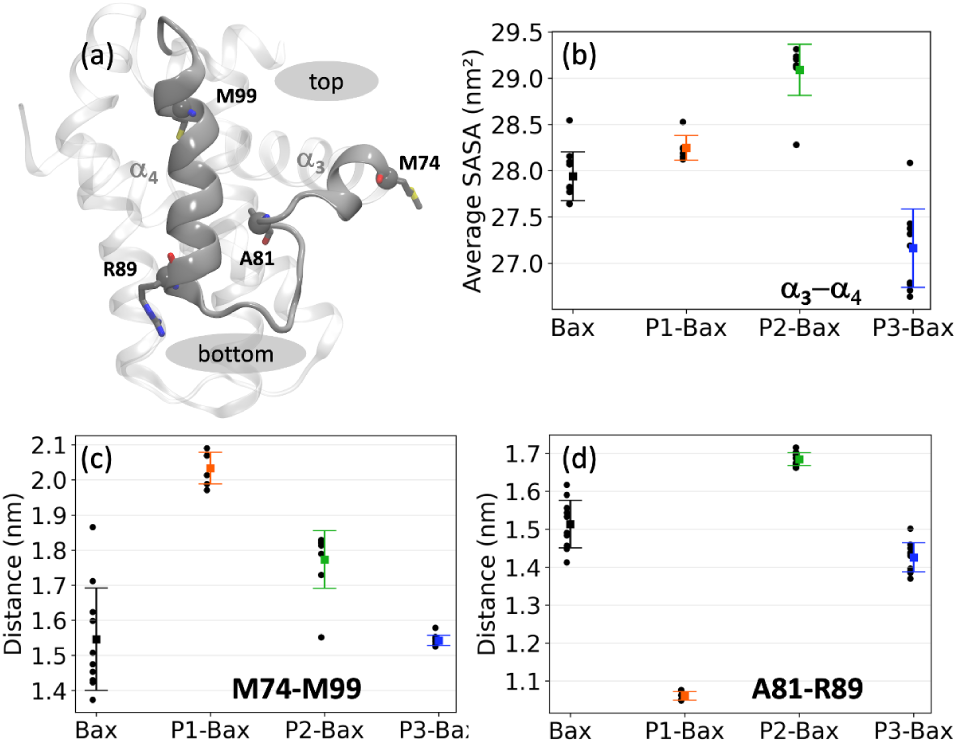
(a) Snapshot of the hydrophobic groove. Highlighted are helices *α*_3_ and *α*_4_ and the the residues used to characterize the *top* and *bottom* sites. (b) Average solvent accessible areas of the hydrophobic groove defined as the M74-M99 sequence encompassing the *α*_3_-*α*_4_ helices. (c) Average distance between the C*_α_* atoms of residues M74 and M99 defining the *top* of the groove. (d) Average distance between the C*_α_* atoms of residues A81 - R89 defining the *bottom* of the groove.

### Intermolecular hydrogen bonds ensure complex stability

The stability of the protein-peptide interface is driven by a series of hydrogen bonds. These vary with protein sequence ranging from 2.1 ± 0.3 upon P1 binding to 3.8 ± 0.3 and 5.1 ± 0.4 for P2 and P3, respectively. P1 binds on the surface of the canonical groove at the BH3 domain contacting the *α*_2_-helix and the C-terminus of *α*_8_. The peptide is “encapsulated” around the side chain of residue K58 with two major stabilizing contacts being hydrogen bond pairs K58-E44 and K58-G38. A quantitative analysis revealed that the backbone of the cyclic peptide interacts with the K58 side chain over 90%, of the simulation time (Fig. 4(a)). A series of other H-bonds and salt bridges were identified to contribute to a lesser degree to the stability of the complex. The P1 binding site is shared by vMIA, a previously identified Bax suppressor. ^16^ The mechanism of action of the suppressor is to position itself in such a way to simultaneously stabilise the *α*_3_-*α*_4_ and *α*_5_-*α*_6_ hairpins, preventing the conformational changes that Bax needs to undergo for its mitochondrial outer membrane insertion and oligomerization, events that invariably lead to the death of the cell.

**Figure 4:**
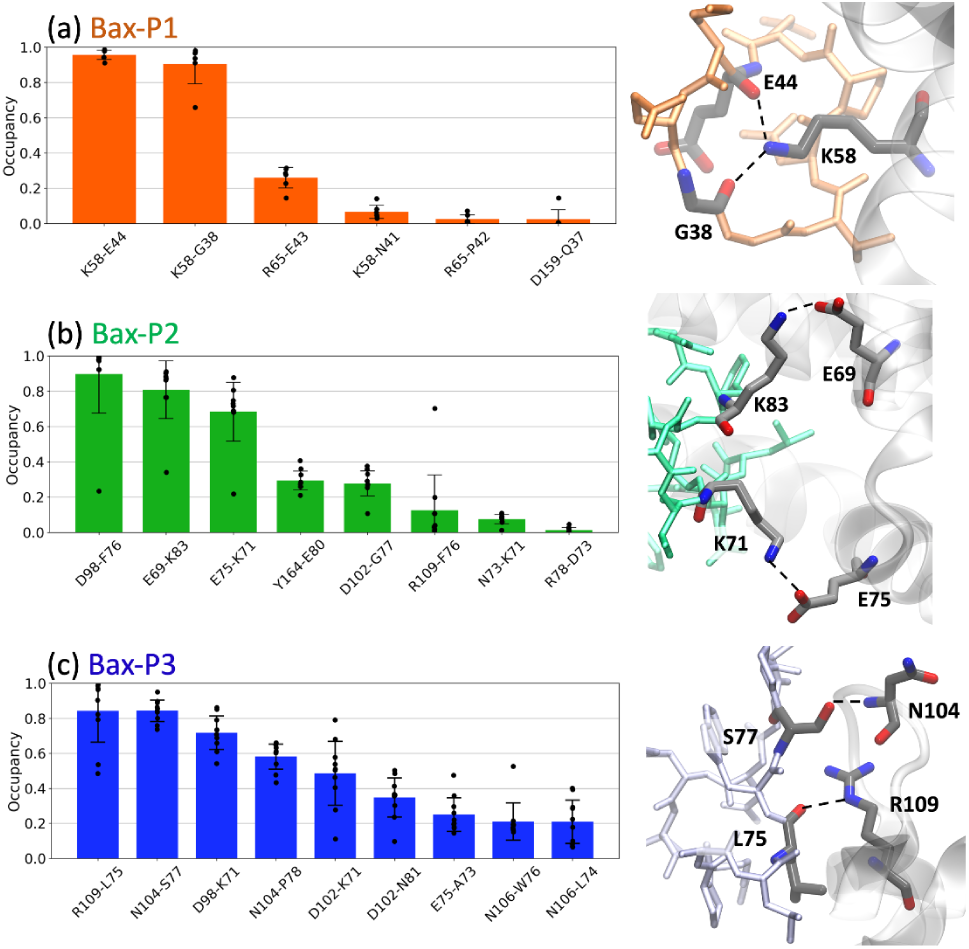
Hydrogen bond between Bax and the individual peptides. A hydrogen bond is considered if the donor-H acceptor distance *<*0.25 nm and for the donor-H acceptor angle *>*120° The error bars represent the standard error of the mean calculated as the standard deviation of the average values over the independent runs (black points).

Peptides P2 and P3 insert into the canonical hydrophobic cleft of Bax, which is the binding target of many BH3 only *activator* proteins such as Bid, Bim, and Bad. ^50–52^ The peptides mimic key hydrophobic and polar contacts of the BH3 ligands. Three major contacts contribute the most to the stability of the P2-Bax complex, *i.e.,* the backbone F76-nitrogen D98 H-bond, and the two E69-K83 and E75-K71 salt bridges. The corroborated effect of the less populated D102-G77 and R109-F76 interactions, which encompass residues on the *α*_4_ helix and the *α*_4_-*α*_5_ loop, and the F76-D98 hydrogen bond is the reduced flexibility of the *α*_4_ helix and the subsequent loop. The less frequent interaction between between Bax Y164 and peptide E80 may contribute to the stabilization of the C-terminus of Bax. Despite its shorter and distinct sequence as compared to P2 (13 residues as opposed to the 16 residues), P3 forms more contacts with Bax and the two peptides both interact with residues D98, E75, D102 and R109 in Bax. The P3-Bax interface is largely stabilized by five interactions (N104-S77, N104-P78, R109-L75, D98-K71 and D102-K71), which are situated at the *α*_4_ and *α*_5_ helices (Fig. 4(c)). The long-lived R109-L75 salt bridge and N104-S77 hydrogen bond located at the *α*_4_-*α*_5_ loop may contribute to the reduced flexibility observed in the RMSF (Fig. 2(b)).

### Bax stabilizes specific peptide conformations

To investigate the effects of Bax on the conformations of the peptides, each peptide was individually subjected to ten 500-ns simulations in absence of the protein. The starting conformations of the free peptides were extracted from the simulations of the complexes, *i.e.,* the initial conformation corresponds to a stably bound peptide conformation (see Methods). The peptides in the free state sample conformations that deviate from the bound configurations, with C*_α_* RMSD values relative to their bound states of 0.16 ± 0.01 nm for P1, 0.24 ± 0.07 nm for P2 and 0.24 ± 0.02 nm (Fig. S3). The analysis focused on the P1 intramolecular contacts revealed that the free peptide maintains the internal E44-N41 H-bond (Fig. 5(a)), yet to lesser extent as compared to the complexed state. A series of distinct contacts are transiently formed in the free state, which indicates that the peptide is more dynamic than in the bound conformation. Additionally, in both the bound and unbound state, the peptide is devoid of any secondary structure and is rich in coils and turns (Fig. S5(a)).

**Figure 5:**
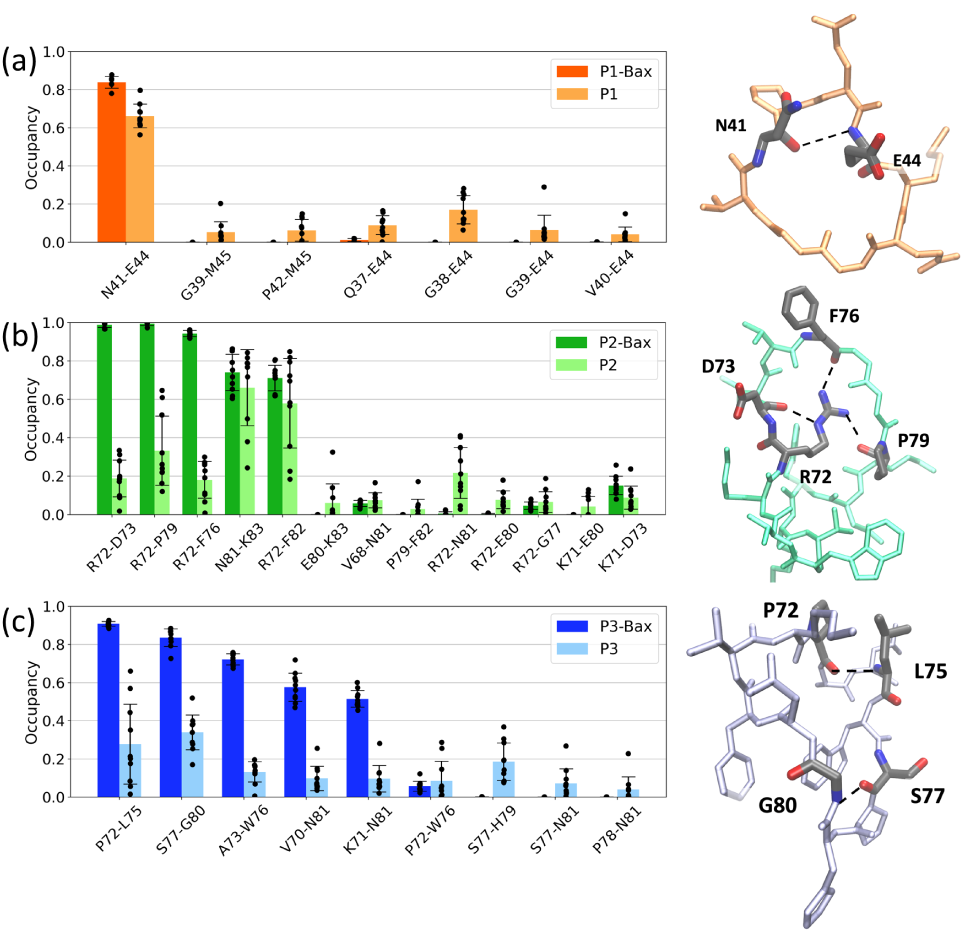
Peptide intramolecular hydrogen bond occupancy. Compared are the contacts in the bound state (dark color) and the unbound state (light color, left panels). Shown are representative snapshots of the peptides in the bound state with the residues involved in the most populated contacts highlighted (right snapshots). A hydrogen bond is considered if the donor-H acceptor distance *<*0.25 nm and for the donor-H acceptor angle *>*120°. The error bars represent the standard error of the mean calculated as the standard deviation of the average values over the independent runs (black points).

In contrast, the differences between the bound and unbound state of P2 and P3 are more pronounced. In complex, P2 attains a twisted boat conformation, rationalized via “reduction” to cycloalkane conformations, where the side chain of R72 is inserted between the peptide backbones, hence contributing to the stability of the peptide. The conformation is lost in the unbound simulations as the peptide moves from a boat conformation to a chair conformation where the side chain of R72 is no longer engaged in stabilizing interactions with the peptide backbone. Instead, the hydrophobic phenyl group of F82 occasionally moves to maintain the bound state hydrogen bonds K83-N81 and F82-R72. Nevertheless, the dynamics of these contacts reflects in the larger error bars as compared to the bound state (Fig. 5(b)). Thus, the protein favors the twisted boat conformation, which is sampled to a lesser extent in the free simulations (Fig. S7). As for P1, P2 is devoid of secondary structure elements and is rich in coils and turns (Fig. S5(b)).

Similar to P2, the predominant intramolecular interactions in the bound P3 are significantly reduced in the free state. The peptide in complex with Bax, is stabilized by five hydrogen bonds. Furthermore, in the bound state, the A73-L75 segment predominantly adopts a 3_10_ helical conformation, a structure marginally sampled in the unbound simulations (Fig. S6). The free peptide is dynamic as highlighted by the large error bars in the contact occupancies (Fig. 5(c)) and devoid of secondary structure elements (Fig. S5(c)).

### Favorable peptide binding

Having characterized the structural details of the protein, the complexes and the peptides independently, the next natural step is to quantify the binding of the peptides to Bax. For this, four sets of umbrella sampling simulations were performed, one for each peptide and one for the negative control (see Methods). Because the peptides bind at different locations and have different binding modes, the profiles in Fig. 6 were shifted along the reaction coordinate for ease of comparison. This does not affect the calculation of the binding free energies. The negative control peptide was constructed from the sequence P1 to disrupt hydrogen bond formation and/or induce steric clashes (see Methods).

**Figure 6:**
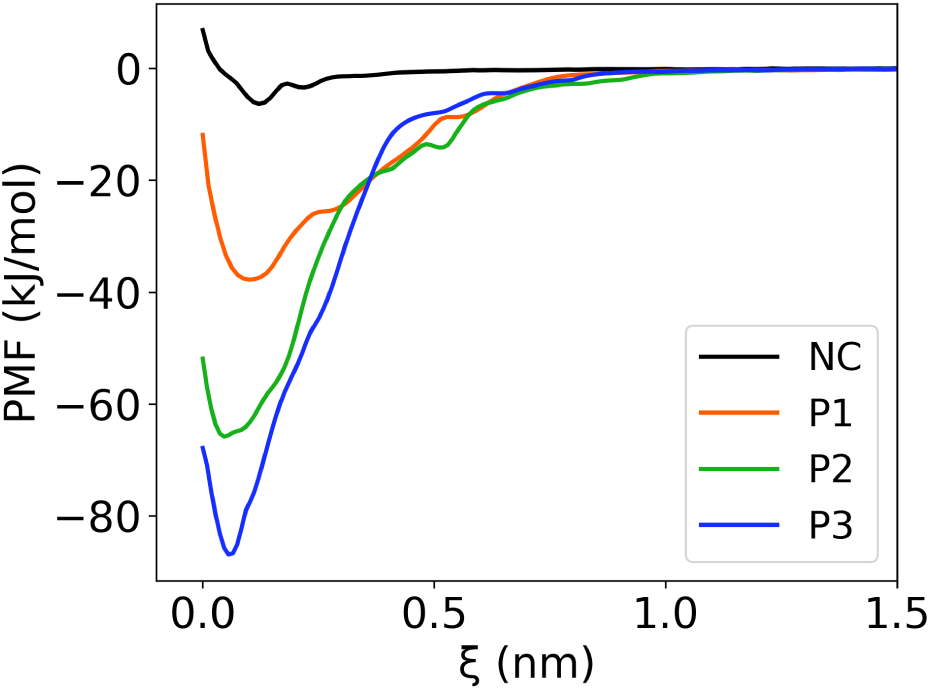
Free energy curves as a function of the center of mass distances between Bax and the peptides. The black curve corresponds to the profile of the negative control (NC).

The free energy curves of all peptides show a minimum at the optimal attachment sites of the peptides, which corresponds to the center of mass distance between the protein and the peptide (Fig. 6). The negative control has a relatively shallow minimum as compared to the designed peptides. This stands as evidence for the lack of stabilizing contacts in the complex and is in line with the conventional molecular dynamics simulations, which show no long-lived attachment of NC to Bax. Peptides P1, P2 and P3 all remained in stable contact during the MD simulations. In the P1-Bax complex, the two hydrogen bonds (K58-E44 and K58-G38) are preserved at the minimum of the free energy profile. Upon pulling the peptide away from the protein, the hydrogen bonds start alternating between the two residues until no contact is established at longer distances. The gradual breaking of the hydrogen bonds reflects in a broad free energy curve (Fig. 6). The profile of the P2-Bax complex presents a lower minimum, which is ascribed to four stabilizing hydrogen bonds (D98-F76, E69-K83, E75-K71 and D102-G77). Similar as in the case of P1, the detachment is gradual and occurs with the step wise breaking of the H-bonds, *i.e.,* D98-F76 is the first bond to break, followed by E69-K83 and N73-K71, and subsequently D102-KG77. The P3-Bax profile shows the lowest minimum across the tested peptides, which corresponds to five dominant hydrogen bonds (R109-L75, N104-S77, D98-K71, N104-P78, D102-K71). Upon pulling the peptide away from the protein, four bonds break simultaneously (N104-S77, N104-P78, R109-L75, and D102-N81), which results in a steeper profile as compared to P1 and P2. The last contact to break is the D102-K71 salt bridge.

To estimate the binding free energies, the detachment free energies of each peptide are first calculated by^53,54^

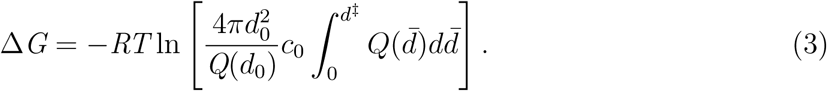

Here, the partition function as a function of distance, *Q*(*d*), is derived from simulations for short distances using *Q*(*d*) = exp[−*A*(*d*)*/*(*RT*)]. For distances beyond the interaction range, an entropy-dominated extrapolation is used: 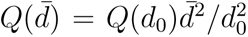. Here, *d*_0_ is an arbitrary distance beyond the interaction range that merges the two partial solutions, *d*^‡^ represents the location of the transition state of the binding reaction and was chosen to be 1.5 nm, *R* is the gas constant, and *c*_0_ is a reference concentration, typically 1 M. The numerical results of the binding free energies are shown in Table 1. These have been evaluated also relative to the negative control. The results show that the number of hydrogen bonds stabilizing the protein-peptide complex contribute the most to the binding free energies. Hence, the binding of a peptide becomes more favorable with increasing number of stabilizing H-bonds as in the specific case of P3.

**Table 1:** Free energy change upon binding to Bax (Δ*G*) and relative to the negative control (ΔΔ*G*).

| Peptide | $\Delta G$ (kJ/mol) | $\Delta\Delta G$ (kJ/mol) |
| --- | --- | --- |
| NC | -7.5 | - |
| P1 | -42 | -34.5 |
| P2 | -70 | -62.5 |
| P3 | -89 | -81.5 |

## Discussion and Conclusion

Neurodegeneration is the leading cause of brain disorders worldwide, and Parkinson’s disease is the fastest growing among them.^55^ Parkinson’s disease involves the loss of dopamine-producing neurons in the brain, leading to motor symptoms. Here, we introduce a novel computational strategy to develop and validate cyclic peptides with the ultimate intent to prevent the initiation of intrinsic apoptosis and possibly prevent or delay the degeneration of neurons as observed in Parkinson’s disease. Our strategy relies on three steps prior to experimental testing. We first rationally design a series of peptides starting from high resolution structures of protein complexes. Subsequently, we computationally validate and optimize the binding of the three peptides to Bax by introducing single point mutations and relying on results from (enhanced sampling) molecular dynamics simulations. Finally, we determine their binding free energies and advance the peptides to experimental testing. Our results reveal that the Bax-peptide complexes are structurally stable on timescales of 5 *µ*s and are stabilized by a series of inter- and intra-molecular hydrogen bonds. Furthermore, the peptides reduced the intrinsic flexibility of the protein by binding at the hydrophobic groove (the dimerization site of Bax^13^) and at allosteric sites. Interestingly, the peptides modulate the dynamics of the canonical hydrophobic groove in a different way. Peptide 3 (^70^VKPALLWSPHGNF^82^) has a marginal impact, while peptide 1 (^37^QGGVNPEEM^45^) and peptide 2 (^68^VWVKRDLVFGGPENFK^83^) both open the hydrophobic groove. The latter has a more pronounced impact, which can potentially impede the ability of Bax to oligomerize, a hypothesis we will test experimentally. Finally, we determined the binding free energies of the peptides to Bax, highlighting the role of the hydrogen bonds and peptide conformational stability in the integrity of the complex.

It has been proposed that Bax activation may require cooperation among various binding sites.^56^ This suggests that finding binders that engage sites beyond the typical hydrophobic groove might lead to allosteric inhibitors. Here, we developed P1, which binds the D53-R65 segment outside the hydrophobic groove but still modulates its dynamics. Hence, further exploration of the impact of one or multiple P1 peptides on Bax, can provide new avenues towards the allosteric modulation of Bax. To access the allosteric pocket one might consider enriching the solvent with small molecules that can gently open the pocket without disrupting its secondary structure.^57^

Our results show that P2 and P3 bind with high affinity and at similar interaction sites to Bax. They insert themselves in the so-called S184 cryptic pocket, referring to a pocket near residue S184 on the *α*_9_-helix that is prone to phosphorylation.^58^ Phosphorylation of the S184 residue causes full length Bax to lose its pro-apoptotic function. ^58–60^ The pocket consists of residues at the *α*_4_-*α*_5_ loop and the *α*_9_-helix. P3 binds with stronger affinity than P2, which is ascribed to more intermolecular contacts with Bax on the *α*_4_-*α*_5_ helices and the connecting loop. Hence this peptide would arguably fit better in the cryptic S184 pocket than peptide P2. Binding at that site mimics pro-survival proteins and may thus result in inhibition of Bax. However, Bax activators such as the *α*-helical BidBH3 peptide, also occupy the canonical hydrophobic surface groove. ^61^ Notably, upon binding of these activators, cavities appear on Bax at the interface between the core and latch domains (*α*_2_, *α*_5_ and *α*_8_).^13,62^ Such cavities are destabilizing and suggest that the binders thereby induce conformational changes which lead to release of the core and its *α*_2_ segment.^63^ These cavities do not form in complexes with pro-survival proteins^13^ and are not accessible on the timescales of our simulations.

As mentioned in the introduction, our long-term goal is to create a machine learning powered platform for peptide design.^19^ This platform would work as a digital twin and rely on input from computational and experimental results to generate novel and better binders towards a target. Evidently, building such a platform is not effortless. As such, the present study sets the stage for generating the computational component of the platform. Future studies will address further computational optimization and experimental validation of the developed peptides. Importantly, the concepts and strategies introduced here extend beyond drug design and will aid in the development of novel bio-inspired materials.

In conclusion, this study lays the foundation for the iterative development of peptides that bind to Bax. We introduced and validated a new protocol for digital peptide development, in which we rationally designed and computationally dissected three cyclic peptides, which can modulate the dynamics of the canonical hydrophobic groove (Bax dimerization site^13^). Furthermore, the stability of a Bax-peptide complex is maintained by a series of inter- and intra-molecular hydrogen bonds, which reflects in the calculated binding free energies. Importantly, this novel protocol can be easily tailored and extended to other protein targets, for which peptides represent attractive binders and for which suitable high resolution structures and/or dynamics studies are available. ^64,65^

## Acknowledgement

I.M.I. acknowledges support from the Sectorplan Bèta & Techniek of the Dutch Government and the Dementia Research - Synapsis Foundation Switzerland. I.M.I. and L.H. acknowledge support from the Molecular Material Design Technology Impulse grant.

