## Supplementary Information for "Computational design of Bax-inhibiting peptides"

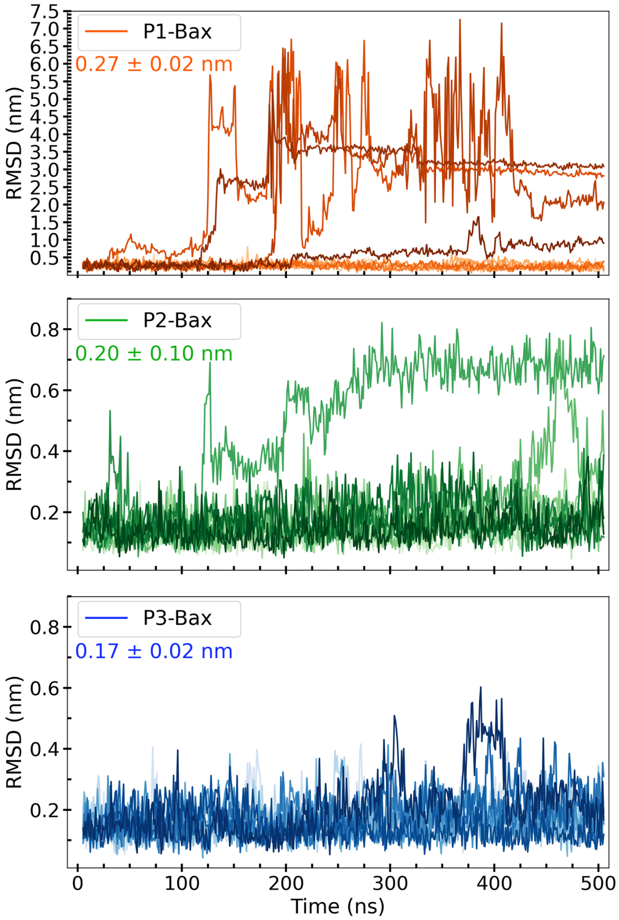


Fig. S1: **Peptide stability.** Shown are the time series of the displacement of the peptides with respect to Bax. First the structural alignment of the individual snapshots saved along the MD simulations is carried out on the $C_\alpha$ atoms of the secondary structure elements of Bax. Then for each MD snapshot the peptide $C_\alpha$ root-mean-square deviation (RMSD) is calculated as $\sqrt{\frac{1}{N_{ab}}\sum_{i=1}^{N_{ab}}(\mathbf{r_i}-\mathbf{r_i^{ref}})^\mathbf{2}}$, where $\mathbf{r_i}$ and $\mathbf{r_i^{ref}}$ are the actual and reference coordinates, respectively, of the peptide $C_\alpha$ atom $i$. $N_{ab}$ is the number of residues in the peptide. The high RMSD values for three of the ten P1 complexes indicate that the peptide detached from the Bax surface. Hence, for the analysis only the stable complexes are used. The values in the top left corners of the plots represent the average RMSDs and the errors represent the standard error of the mean calculated as the standard deviation of the average values over the independent stable runs.

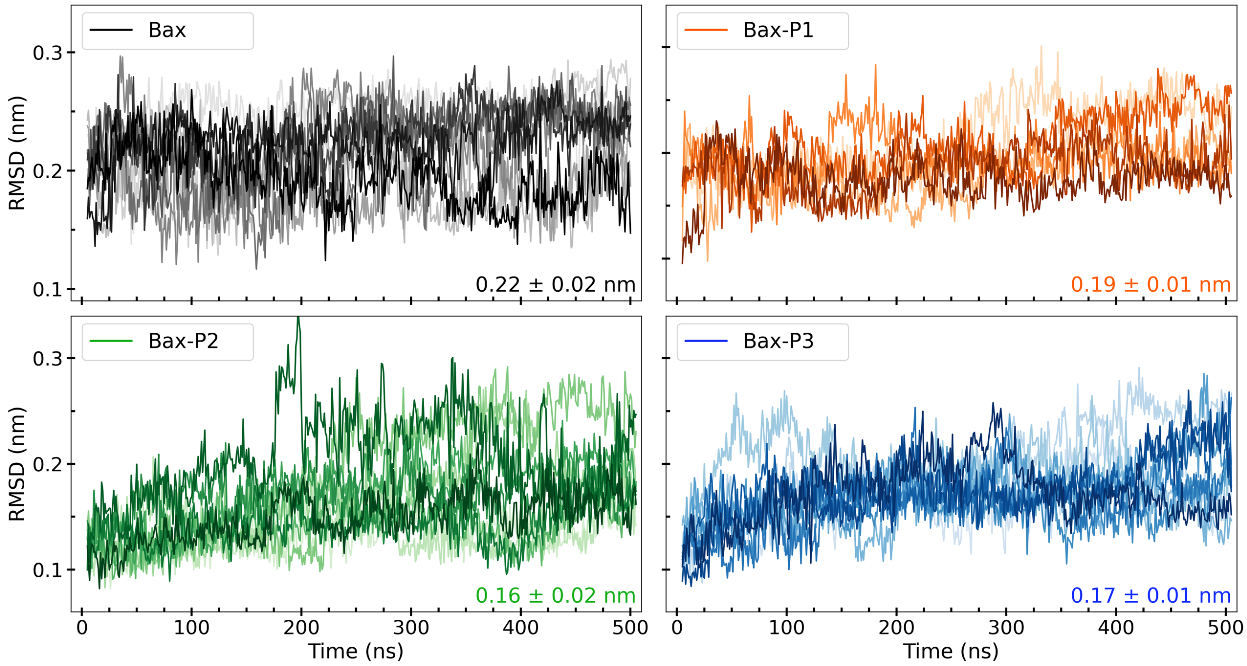


Fig. S2: **Protein stability.** Shown are the time series of the displacement of Bax with respect to the crystal structure. First the structural alignment of the individual snapshots saved along the MD simulations is carried out on the $C_\alpha$ atoms of the secondary structure elements of Bax. Then for each MD snapshot the peptide $C_\alpha$ root-mean-square deviation (RMSD) is calculated as $\sqrt{\frac{1}{N_{ab}}\sum_{i=1}^{N_{ab}}(\mathbf{r_i}-\mathbf{r_i^{ref}})^2}$, where $\mathbf{r_i}$ and $\mathbf{r_i^{ref}}$ are the actual and reference coordinates, respectively, of the protein $C_\alpha$ atom $i$. $N_{ab}$ is the number of residues in Bax. The values in the lower right corners of the plots represent the average RMSDs and the errors represent the standard error of the mean calculated as the standard deviation of the average values over the independent runs (in which the peptides remain stably attached to Bax).

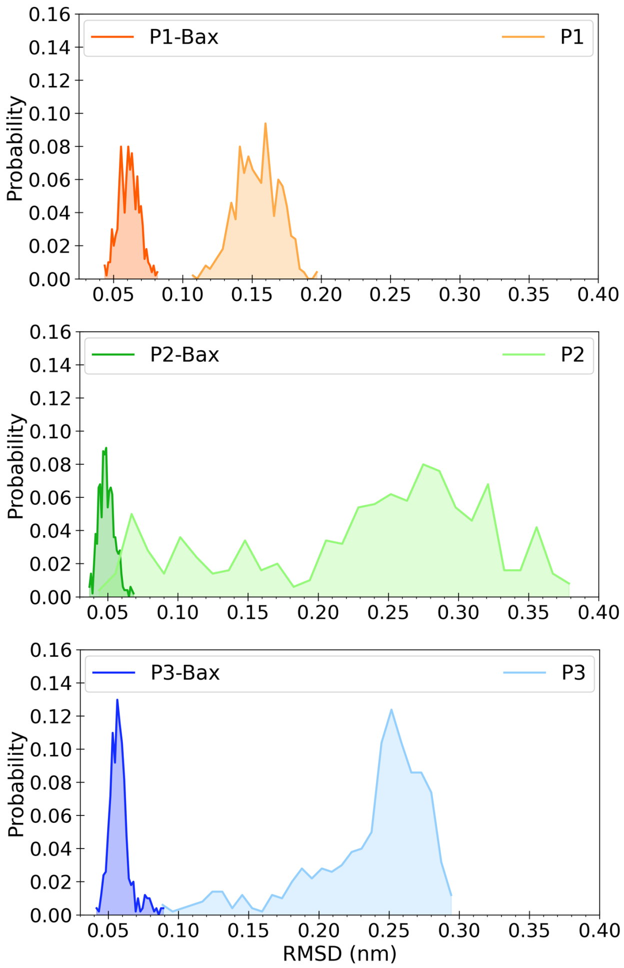


Fig. S3: **Peptide conformations.** Shown are the probability distributions of the peptide RMSDs of the in the bound (dark colors) and unbound states (light colors) relative to the bound state. The unbound peptide runs are started from the bound peptide conformations in absence of Bax. The analysis shows that the unbound peptides sample states that are different from the bound conformations, suggesting that Bax stabilizes specific intramolecular peptide bonds.

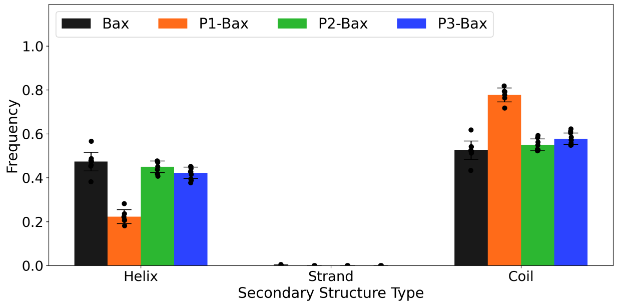


Fig. S4: **Secondary structure of $\alpha_3$.** Shown are the normalized secondary structure assignments of $\alpha_3$ and the $\alpha_3$-$\alpha_4$ loop in absence and presence of the peptides.

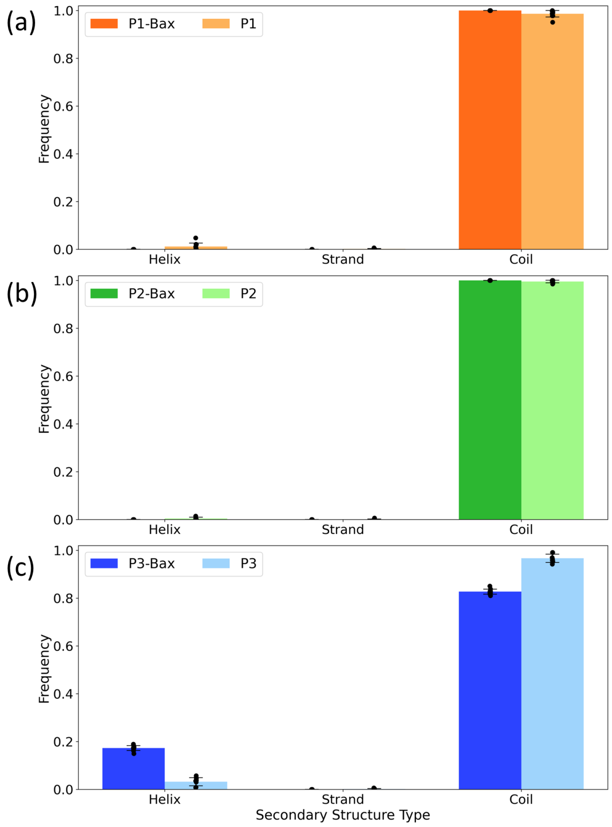


Fig. S5: **Peptide secondary structure assignment.** Shown are the normalized secondary structure assignments of the peptides in the bound and unbound states. The results show that the peptides adopt predominantly disordered structures, except for P3, which partially adopts and $\alpha$-helical structure when bound to Bax.

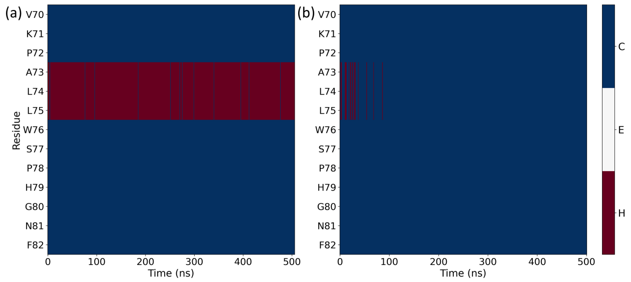


Fig. S6: **Secondary structure of P3.** Shown are the secondary structure time evolutions of P3 for the (a) bound and (b) unbound states, with C = coil, E = strand and H = helix.

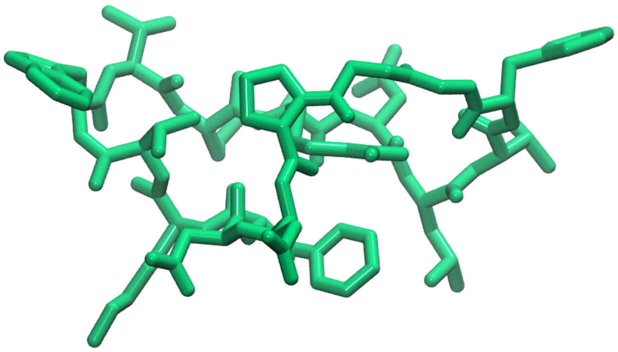

Fig. S7: **P2 twisted boat conformation** Shown is the twisted boat conformation of peptide P2.
